# Identification and structural modeling of the chlamydial RNA polymerase omega subunit

**DOI:** 10.1101/2022.09.22.509108

**Authors:** Andrew Cheng, Danny Wan, Arkaprabha Ghatak, Chengyuan Wang, Deyu Feng, Joseph D. Fondell, Richard H. Ebright, Huizhou Fan

## Abstract

Gene transcription in bacteria is carried out by the multisubunit RNA polymerase (RNAP), which is composed of a catalytic core enzyme and a promoter-recognizing σ factor. RNAP core enzyme comprises two α subunits, one β subunit, one β’
s subunit, and one ω (omega) subunit. Across multiple bacterial taxa, the RNAP ω subunit plays critical roles in the assembly of RNAP core enzyme and in other cellular functions, including regulation of bacterial growth, stress response, and biofilm formation. However, for several intracellular bacterium, including the obligate intracellular bacterium *Chlamydia*, no RNAP ω subunit previously has been identified. Here, we report the identification of *Chlamydia trachomatis* hypothetical protein CTL0286 as the chlamydial RNAP ω ortholog, based on sequence, synteny, and AlphaFold and AlphaFold-Multimer three-dimensional-structure predictions. We conclude that CTL0286 functions as the previously missing chlamydial ω ortholog. Extensions of our analysis indicate that all obligate intracellular bacteria have ω orthologs.

**IMPORTANCE:** Chlamydiae are common mammalian pathogens. Chlamydiae have a unique developmental cycle characterized with an infectious but nondividing elementary body (EB), which can temporarily survive outside host cells, and a noninfectious reticulate body (RB), which replicates only intracellularly. Chlamydial development inside host cells can be arrested during persistence in response to adverse environmental conditions. Transcription plays a central role in the progression of the chlamydial developmental cycle as well as entry into and recovery from persistence. The identification of the elusive ω subunit of chlamydial RNAP makes possible future study of its regulatory roles in gene expression during chlamydial growth, development, and stress responses. This discovery also paves the way to prepare and study the intact chlamydial RNAP and its interactions with inhibitors *in vitro*.

## INTRODUCTION

RNA synthesis in bacteria is carried by a single RNA polymerase (RNAP). The bacterial RNAP is a multisubunit enzyme (1). In almost all bacteria, the catalytic core enzyme of the RNAP (RNAP core) is composed of two α subunits, one β subunit, one β’ subunit, and one ω subunit (1, 2). Association of a σ factor to the core enzyme results in the formation of the RNAP holoenzyme (1). In the context of the holoenzyme, the σ factor is the primary determinant of promoter recognition and binding, and the RNAP core catalyzes the initiation and elongation of RNA synthesis using DNA as template (2-4).

The RNAP ω subunit, a protein of only about 10 kDa, initially was thought to be a contaminant in purified RNAP preparations (5-7). This view was prompted by the observation that ω-free RNAP preparations were active in transcription assays (8). However, the observation of increased transcription-initiation activity by an RNAP derivatives having ω fused to DNA-binding domains indicated ω was an integral component of RNAP (9). Further studies showed that ω is critical for the folding of the RNAP β’ subunit and the for the assembly and stability of RNAP core enzyme (10-14). Studies using ω*-*deficient bacteria showed that ω is important for response to amino acid starvation, thermal and CO_2_ acclimation, biofilm formation, and antibiotic production, and also affects growth under standard culture conditions (15-20). It was also shown that ω regulates the association of principal and alternative σ factors by the RNAP core enzyme and thus can affect promoter-recognition selectivity (21-23). Taken together, these and other studies suggest that ω serves as an important component of the bacterial RNAP holoenzyme and is required for numerous physiological functions [for review, see (24-26)]. ω has been found in all free-living bacteria and in some obligate intracellular bacteria (24, 25). ω also is present in some eukaryotic chloroplasts (27). An ortholog of ω, termed RpoK, is present in archaeal RNAP (28), and an ortholog of ω, termed RPB6, is present in eukaryotic RNAP I, II, and III (29).

Chlamydiae are intracellular bacteria that replicate only inside eukaryotic host cells (30, 31). Chlamydiae and *Chlamydia-*like organisms have been isolated from a wide range of hosts (32-46). Significantly, *Chlamydia trachomatis* is the number one sexually transmitted bacterial pathogen globally, and also is a major cause of preventable blindness in developing countries (47-49), and *C. pneumoniae* is a common respiratory pathogen (50-54). Several animal *Chlamydia* species are zoonotic pathogens (55-64). *Waddlia chondrophila* is one of several *Chlamydia*-like organisms, termed environmental chlamydiae, typically found in lower eukaryotes, such as amoebae, *but* can infect, and induce abortion in, vertebrates, including humans (65).

Chlamydiae are characterized by a unique developmental cycle consisting of two distinct cellular forms. The infectious but non-proliferative elementary body (EB) is capable of temporarily surviving in extracellular environments and invading host cells. Following invasion of host cells and entry into cytoplasmic vacuoles, EBs differentiate into proliferative reticulate bodies (RBs). Following multiple rounds of replication, RBs convert back into EBs, which then exit host cells (66-68). In addition to this “productive” chlamydial developmental cycle, under unfavorable environmental conditions (e.g., nutrient/mineral starvation, increased temperature, or exposure to inhibitory antibiotics, or cytokines), chlamydiae can enter into a “persistent” state characterized by aberrant RBs inside infected cells, and, when environmental conditions improve, the aberrant RBs can exit the persistent state and resume production of EBs, (69-75).

Both the productive chlamydial developmental cycle and persistent infection are controlled by gene transcription (69, 71, 75-77). The chlamydial genome encodes three σ factors (σ^66^, σ^28^ and σ^54^), as well as the α, β and β’
s subunits of the core enzyme (78, 79). Surprisingly, it previously has not been possible to identify a candidate gene encoding the ω subunit in any chlamydial genome [e.g., (80-83)]. In principle, the chlamydial *rpoZ* gene may have been lost in the evolutionary process during which *Chlamydia* reduced its genome size to adapt to its unique developmental cycle. Alternatively, in principle, the chlamydial ω protein may have gone undetected due to low sequence homology with known bacterial and chloroplast ω factors.

Here, we report the identification of chlamydial ω, based on conserved amino-acid sequence, conserved synteny, and AlphaFold-predicted conserved three-dimensional structure and interactions. In addition, we also present an AlphaFold-Multimer model of the three-dimensional structure of a complex composed of the chlamydial RNAP β, β’, and ω subunits. The identification of the previously elusive chlamydial ω sets the stage for investigation of its roles in regulation of gene expression during chlamydial growth, development, and stress responses. Our findings also set the stage to reconstitute the intact cRNAP from recombinant subunits *in vitro*, for future structural studies and for discovery and development of small-molecule inhibitors as possible anti-chlamydial drugs.

## METHODS

### BlastP analysis

Web-based BlastP was performed at https://blast.ncbi.nlm.nih.gov/Blast.cgi?PAGE=Proteins using default settings (84). Multiple protein sequence alignment was performed with ClustalX2 on a Windows computer on PC (85) or Clustal Omega at https://www.ebi.ac.uk/Tools/msa/clustalo/ using default settings (86, 87).

Three-dimensional-structure prediction AlphaFold version 2.2.0 (88) was installed locally and run using the reduced database option with a maximum template date of November 1, 2021 and the multimer preset enabled for the cRNAP β’-CTL0286 and cRNAP β-β’-CTL0286 complex predictions. The multimer predictions were run with the default pre-trained AlphaFold-Multimer models (89), and the ranked 0 predictions (i.e., with lowest predicted local distance difference test [pLDDT] scores) were used for the figures for each complex. The CTL0286 monomer structure prediction was performed with the default monomer preset and the full database option and used the original CASP14 monomer models without ensembling. Model prediction and amber relaxation were performed for all predictions using a single NVIDIA Tesla V100 Volta GPU with 16GB of memory. Since the total sequence lengths significantly increase the space complexity, forced unified memory was enabled, and the XLA memory fraction environmental variable was set to 4.0 to avoid out of memory errors during runtime.

### Three-dimensional-structure similarity analysis

Structural homology search for AlphaFold model of CTL0286 was performed using the Dali server heuristic PDB Search option (90, 91) available at http://ekhidna2.biocenter.helsinki.fi/dali/. A PDB90 non-redundant subset at 90% sequence identity was used.

### Synteny analysis

CSBFinder-S (v0.6.3) (92) was used with the default settings to find the *gmk-rpoZ* synteny across 23,517 fully sequenced bacterial genomes downloaded from the NCBI genome database. DeepNOG (v1.2.3) (93) was run using the default setting to obtain the COG (Clusters of Orthologous Genes) ID for each gene. Strand information was obtained from the corresponding genomic.gff file for every genome downloaded. The *gmk-rpoZ* synteny was identified by finding COG0194 (*gmk*) and COG1758 (*rpoZ*) together within the CSBFinder-S output.

## RESULTS

### Identification of chlamydial ω: sequence similarity

Although ω has not been detected in Chlamydiae, ω had been detected in two other intracellular bacteria: *Rickettsia* and *Coxiella* (94, 95). Therefore, as a starting point to determine if chlamydiae encode an ω subunit, we performed BlastP analysis for chlamydial genomes using the amino-acid sequences of *R. rickettsii* ω and *C. burnettii* ω (94, 95) as queries. Using default parameters (84), the analysis did not detect sequence homolog to the *R. rickettsii* ω; in chlamydiae. However, the analysis did detect a possible sequence homolog of *C. burnettii* ω: Wcw_0707, a hypothetical protein encoded by the genome of the *Chlamydia-*like organism *W. chondrophila* (82) (Fig. 1A). Analysis of the sequence of Wcw_0707 revealed two features consistent with Wcw_0707 being an ω ortholog. First, Wcw_0707 is 107 amino acids long, similar in size to ω (∼100 amino acids). Second, the Wcw_0707 *N-*terminal region (residues 7-62) exhibits strong sequence similarity to the *C. burnettii* ω (Fig. 1A) and *Escherichia coli* ω *N-* terminal regions (Fig. 1B), which are known to be responsible for binding to the RNAP β’
s subunit and for facilitating the folding of β’ (96). We hypothesized that Wcw_0707 may be the ω subunit in *W. chondrophila*.

**Fig. 1.**
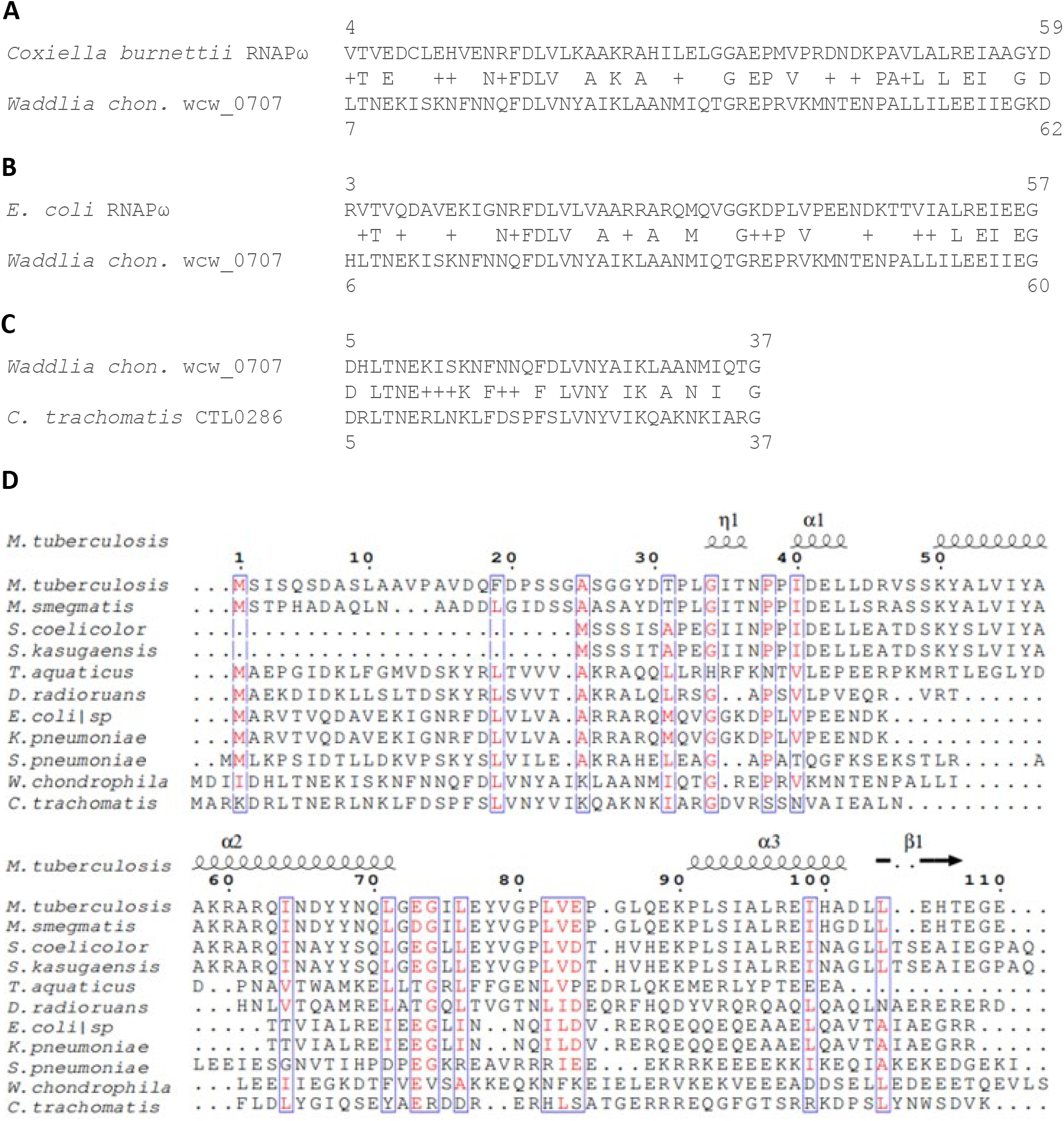
Identification of cRNAP ω candidate by BlastP and sequence alignment. (A) BlastP-detected sequence homology between *Coxiella burnetii* RNAP ω subunit and wcw_0707, a hypothetical protein of the *Chlamydia-*like organism *Waddlia chondrophila*. (B) BlastP-detected sequence homology between *E. coli* RNAP ω and wcw_0707. (C) BlastP-detected sequence homology between wcw_0707 and CTL0286 of *Chlamydia trachomatis*. (D) ClustalX2-detected amino acids conserved in CTL0286 of *C. trachomatis*, wcw_0707 of *W. chondrophila*, and ωs of a variety of bacteria.

Given our primarily interest in transcriptional regulation by the human sexually transmitted pathogen *C. trachomatis*, we next used Wcw_0707 as the query to search for a putative ω gene in the *C. trachomatis* genome. The search revealed a strong sequence similarity between the *N-* terminal region of hypothetical protein CTL0286 of *C. trachomatis* serovar L2 and the N-terminal region of Wcw_0707 of *W. chondrophila* (Fig. 1C). CTL0286 is a small protein of 100 amino acids, similar in length to previously reported RNAP ω subunits and similar in length to Wcw_0707, (81). Notably, although CTL0286 exhibits only low overall sequence similarity to other reported bacterial ω subunits, it contains a key conserved set of amino acids found in ω subunits of a broad range of bacterial taxa (Fig. 1D). Additional BlastP analysis of CTL0286 identified CTL0286 orthologs in all vertebrate chlamydiae (Fig. 2). These findings support the hypothesis that Wcw_0707 is the ω subunit in *W. chondrophila* and enable the hypothesis that CTL0286 and its orthologs are ω subunits in *C. trachomatis* and other vertebrate chlamydiae.

**Fig. 2.**
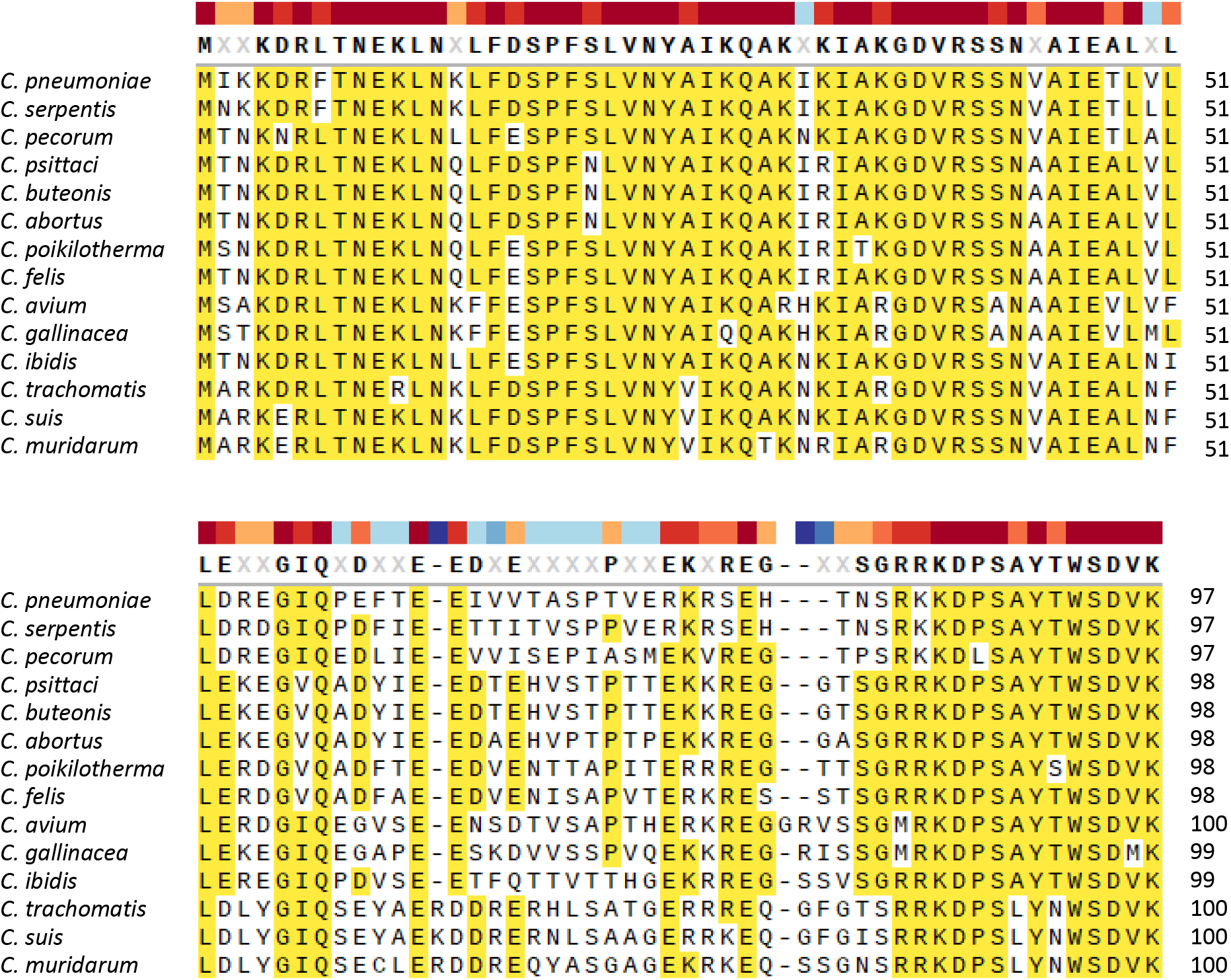
Sequence conservation among ω candidates in all vertebrate chlamydiae. Alignment was performed using ClustalX2.

**Fig. 3.**
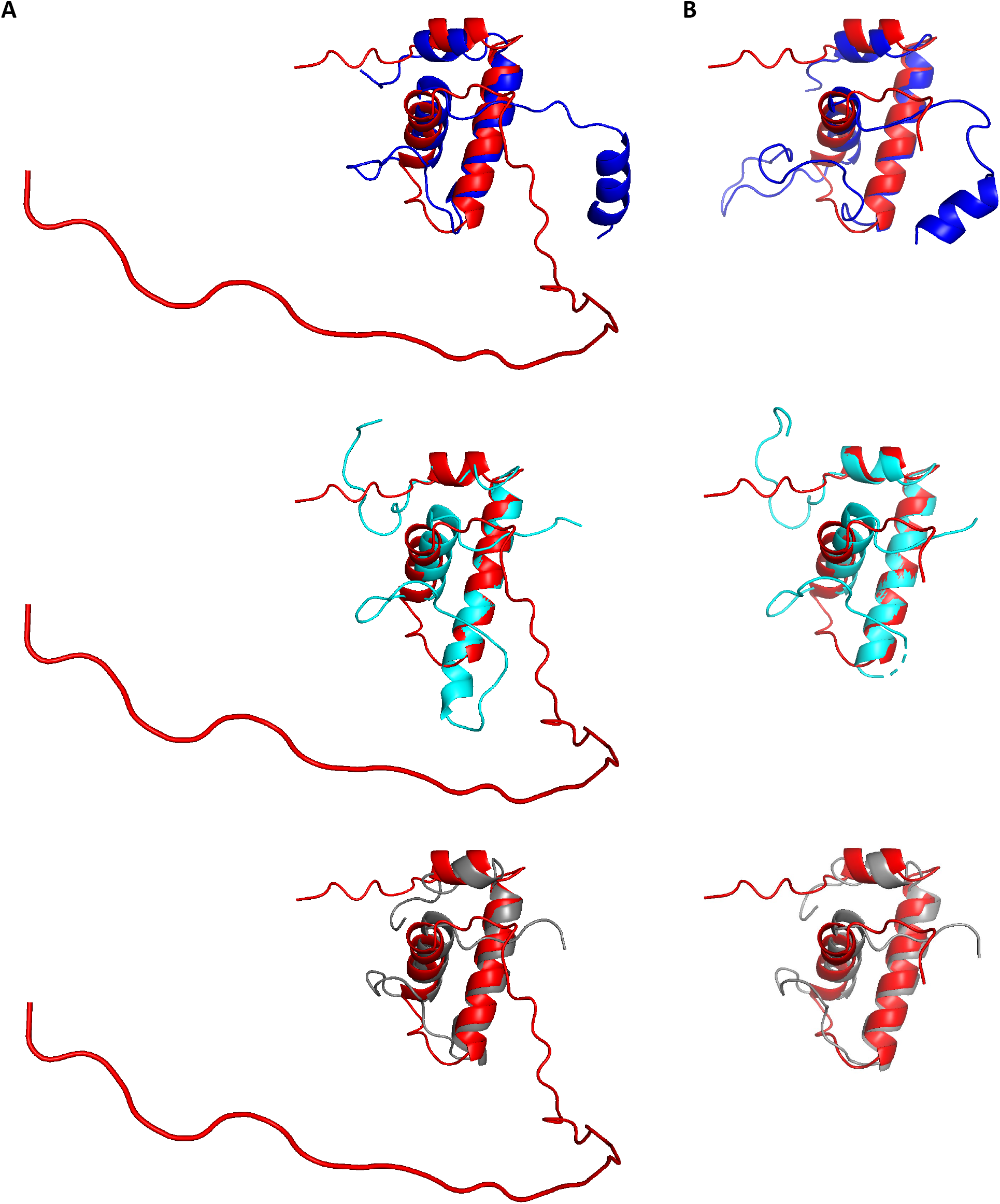
AlphaFold predictions for CTL0286. (A) Superimposition of AlphaFold prediction for full-length CTL0286 (red) on experimental structures of *Clostridium difficile, Mycobacterium tuberculosis*, and *Bacillus subtilis* RNAP ω (blue, cyan, and gray, respectively). (B) Superimposition of AlphaFold prediction for N-terminal region (residues 1-62) of CTL0286 (red) on experimental structures of *Escherichia coli, Mycobacterium tuberculosis*, and *Bacillus subtilis* RNAP ω (blue, cyan, and gray, respectively).

### Identification of chlamydial ω: synteny

Upon manual examination of *rpoZ* in 10 bacterial genomes, we noted that the *rpoZ* gene always is located immediately downstream of the *gmk* gene, which encodes guanylate kinase (Table 1). An *in silico* analysis identified *gmk-rpoZ* synteny in 18302 of 23517 fully-sequenced bacterial genomes. The conservation of *gmk-rpoZ* synteny across a majority of bacteria taxa suggests that there likely is an adaptive advantage to *gmk-rpoZ* synteny, although the character of the adaptive advantage is not readily clear. Interestingly, in *W. chondrophila*, the *wcw_0707* gene is located immediately downstream of the *gmk* gene, and, in all vertebrate chlamydiae species, the *ctl0286* gene and its orthologs are also located immediately downstream of *gmk* (Table 1). This conserved gene order provides further support for the hypothesis that CTL0286 and its orthologs are chlamydial ω subunits.

**Table 1.** Conserved *gmk-rpoZ* linkage in bacterial genomes.

| **Bacterium** | **Gram-stain** | **Upstream gene** | ***rpoZ* or equivalence** | **Downstream gene** |
| --- | --- | --- | --- | --- |
| *Bacillus anthracis* | Positive | *gmk* | *rpoZ* | *coaBC* |
| *Clostridium difficile* | Positive | *gmk* | *rpoZ* | *coaBC* |
| *Lactobacillus acidophilus* | Positive | *gmk* | *rpoZ* | *priA* |
| *Staphylococcus epidermidis* | Positive | *gmk* | *rpoZ* | *SE0887* |
| *Coxiella burnetii* | Negative | *gmk* | *rpoZ* | *spoT* |
| *Escherichia coli* | Negative | *gmk* | *rpoZ* | *spoT* |
| *Haemophilus influenzae* | Negative | *gmk* | *rpoZ* | *spoT* |
| *Vibrio cholerae* | Negative | *gmk* | *rpoZ* | *spoT* |
| *Gardnerella vaginalis* | Variable | *gmk* | *rpoZ* | *dfp* |
| *Mycobacterium tuberculosis* | Variable | *gmk* | *rpoZ* | *metK* |
| ***Waddlia chondrophila*** | Negative | ***gmk*** | ***wcw_0707*** | *wcw_0708* |
| ***Chlamydia trachomatis*** | Negative | ***gmk*** | ***ctl0286*** | *metG* |

### Identification of chlamydial ω: predicted three-dimensional structural similarity

AlphaFold has recently become an indispensable resource for predicting the three-dimensional structures of proteins and protein complexes (88, 89). We first used AlphaFold to predict three-dimensional structure of CTL0286. In the resulting predicted structure for CTL0286, the N-terminal region (residues 1-58) contains three α helices (α1, residues 9-15 residues; α2, residues 19-36; and α3, residues 44-55) that correspond to three α-helices present in all structurally characterized ω subunits (26, 97, 98), and the C-terminal region (residues 58-100) are mostly disordered, similar to in structurally characterized ω subunits having lengths greater ∼60 amino acids (26, 97, 98). Three-dimensional-structure similarity searches of the AlphaFold prediction for full-length CTL0286, performed on the DALI server (90, 91), identified bacterial ω subunits as the three top hits, with Z-scores of 3.8, 3.4, and 3.1, for RNAP ω subunits of *Clostridium difficile* (99), *Mycobacterium tuberculosis* (100), and *Bacillus subtilis* (101), respectively (Table 2). Three-dimensional-structure similarity searches of the AlphaFold prediction for the N-terminal region of CTL0286 (residues 1-62), performed on the DALI server (90, 91), identified bacterial ω subunits as the three top hits, with Z-scores of 5.2, 5.1, and 4.9 for RNAP ω subunits of *Escherichia coli* (102), *Mycobacterium tuberculosis* (103), and *Bacillus subtilis* (104), respectively (Table 2).

**Table 2.** Proteins with structural homology to AlphaFold models of full-length CTL0286 (Fl-CTL0286) or N-terminus of CTL0286 (1-62). Z-score is an optimized similarity score defined as the sum of equivalent residue-wise C α -C α distances among two proteins. Abbreviation: RMSD, Root-mean-square deviation of atomic positions.

| Model | Rank # | PDB structure<br>(reference) | Protein | Bacterium | Z-value | RMSD |
| --- | --- | --- | --- | --- | --- | --- |
| Fl-CTL0286 | 1 | 7L7B (99) | RNAP $\omega$ | *Clostridium difficile* | 3.8 | 3.9 |
| | 2 | 6BZO (100) | RNAP $\omega$ | *Mycobacterium tuberculosis* | 3.4 | 3.1 |
| | 3 | 7CKQ (101) | RNAP $\omega$ | *Bacillus subtilis* | 3.1 | 3.0 |
| CTL0286<br>(1-62) | 1 | 5TJG (102) | RNAP $\omega$ | *Escherichia coli* | 5.2 | 2.1 |
| | 2 | 6KOP (103) | RNAP $\omega$ | *Mycobacterium tuberculosis* | 5.1 | 3.1 |
| | 3 | 7F75 (104) | RNAP $\omega$ | *Bacillus subtilis* | 4.9 | 2.8 |

We next used AlphaFold-Multimer (89) to predict the three-dimensional structure of a complex of CTL0286 and the *C. trachomatis* RNAP β’ subunit (Fig. 4A). The resulting predicted three-dimensional structure of CTL0286-β’ was superimposable, with an rmsd of 2.2 Å for CTL0286 and an rmsd of 4.0 Å for *C. trachomatis* RNAP β’on a crystal structure of the ω-β’ subcomplex of *E. coli* RNAP holoenzyme (PDB 6ALH) (97) (Fig. 4B). Significantly, the predicted three-dimensional structure of CTL0286-β’ includes interactions that bridge the RNAP β’-subunit N- and C-termini (Fig. 4C) as observed in experimental structures of ω-containing RNAP and RNAP complexes (97, 105, 106), where they are believed to reduce configurational entropy of partly folded and folded states of the nearly 1400-residue RNAP β’ subunits, and thereby to facilitate RNAP assembly and enhance RNAP stability (10, 12, 14). We further used AlphaFold-Multimer to predict the three-dimensional structure of a heterotrimeric protein complex comprising CTL0286, *C. trachomatis* RNAP β’, and *C. trachomatis* RNAP β (Fig. 5A). The resulting predicted three-dimensional structure of CTL0286-β’ was superimposable, with rmsd of 2.2 Å for CTL0286 and 2.7 Å for *C. trachomatis* RNAP β, and β, on a crystal structure of the ω-β’-β subcomplex of *E. coli* RNAP holoenzyme (PDB 6ALH) (97) (Fig. 5B) and includes interactions that bridge the N- and C-termini of β’ (Fig. 5C).

**Fig. 4.**
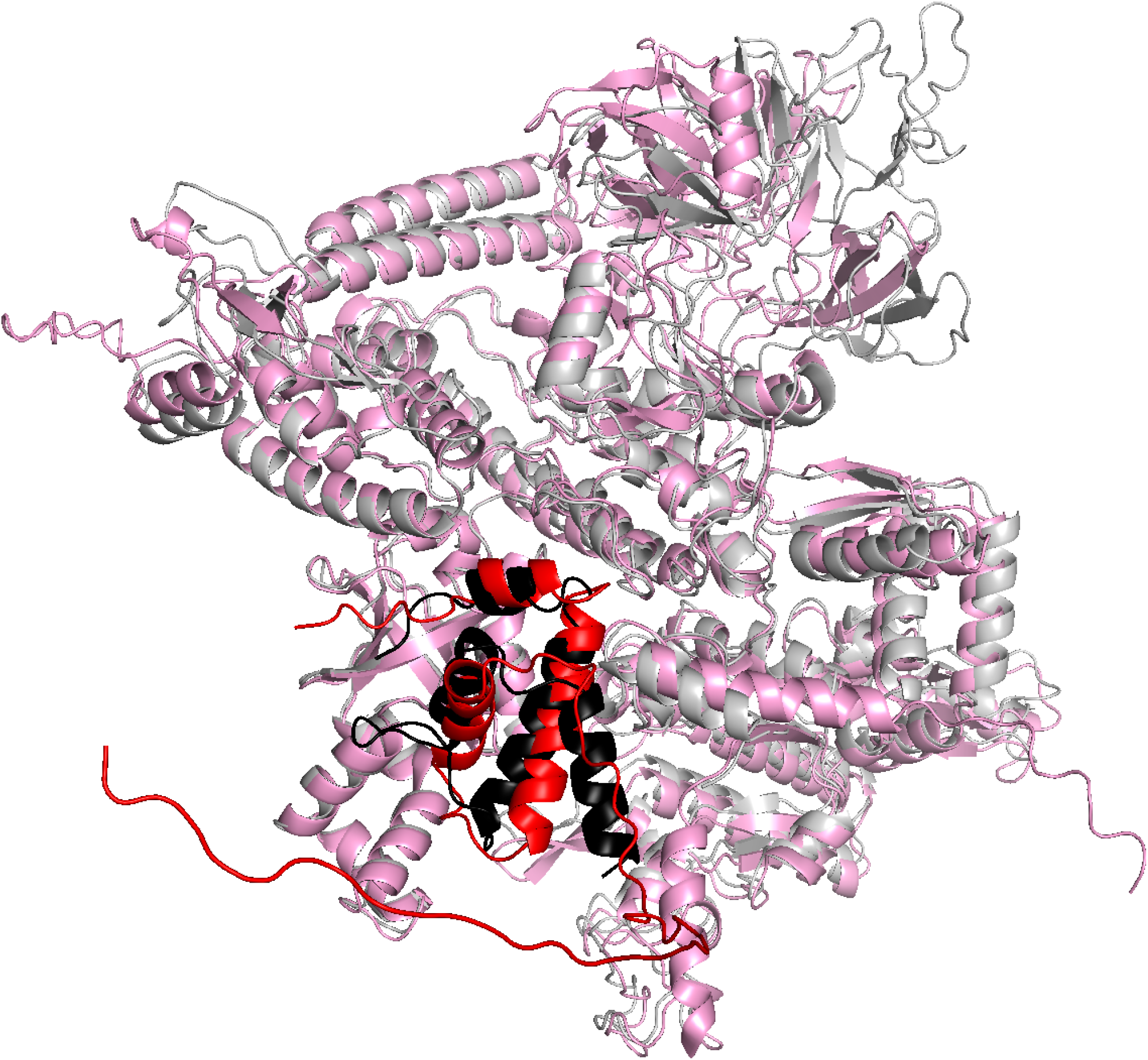
AlphaFold-Multimer predictions for complex comprising CTL0286 and *C. trachomatis* RNAP β’ subunit. Superimposition of AlphaFold-Multimer prediction for CTL0286-β’ (red for CTL0286; pink for β’) on experimental structure of *E. coli* RNAP (PDB 6ALH; black for ω; light gray for β’).

**Fig. 5.**
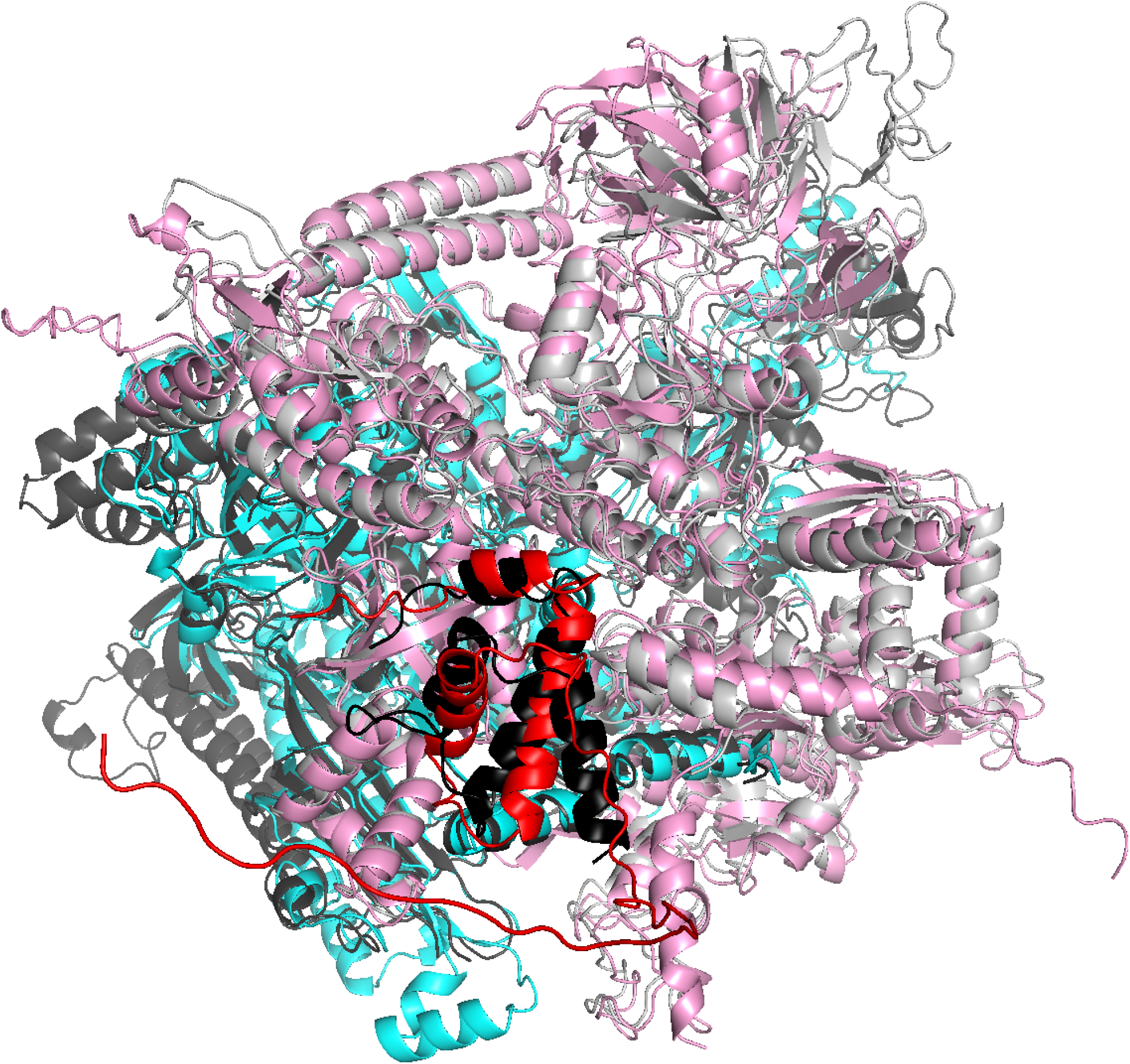
AlphaFold-Multimer predictions for complex comprising CTL0286 and *C. trachomatis* RNAP β’ subunit, and β subunit. Superimposition of AlphaFold-Multimer prediction for CTL0286-β’-β (red for CTL0286; pink for β’; cyan for β) on experimental structure of *E. coli* RNAP (PDB 6ALH; black for ω; light gray for β’; dark gray for β).

Taken together, these findings provide further support for our hypothesis that CTL0286 and its orthologs are *bona fide* chlamydial ω subunits.

### Identification of ω in other obligate intracellular bacteria

After successful identification of RNAP ω subunit in chlamydiae, we next determined if ω is present in other obligate intracellular bacterial taxa beside rickettsiae. NCBI searches identified annotated ω orthologs in the proteomes of *Anaplasma, Ehrlichia, Orientia, Wolbachia* and *Candidatus Midichloria*. Pre-generated AlphaFold structural models of *Anaplasma, Ehrlichia, Orentia*, and *Wolbachia* ω orthologs at www.uniprot.org (107) show three-dimensional strctural similarity to experimentally determined structures of bacterial ω subunits, indicating that the annotations likely are correct. No pre-generated AlphaFold structural model of the annotated *Candidatus Midichloria* ω ortholog is available at www.uniprot.org (107). However, generation of an AlphaFold structural model for the annotated C*andidatus Midichloria* ω ortholog (Fig. 6), followed by three-dimensional-structure similarity searches on the DALI server (90, 91) identified bacterial ω subunits as the three top hits, with Z-scores of 9.3, 8.6, and 8.6 for RNAP ω subunits of *Pseudomonas Aeruginosa* (108), *Mycobacterium tuberculosis* (100), and *Xanthomonos oryzae* (109), respectively, indicating that the annotation likely is correct (Table 3). We conclude that *Analplasma, Ehrlichia, Orientia, Wolbachia*, and *Candidatus Midichloria* all possess RNAP ω subunits.

**Table 3.** Proteins with structural homology to AlphaFold model of annotated *Candidatus Midichloria* RNAP ω.

| Rank # | PDB structure<br>(reference) | Protein | Bacterium | Z-value | RMSD |
| --- | --- | --- | --- | --- | --- |
| 1 | 7XL3 (108) | RNAP ω | *Pseudomonas aeruginosa* | 9.3 | 2.7 |
| 2 | 7L7B (100) | RNAP ω | *Mycobacterium tuberculosis* | 8.6 | 3.0 |
| 3 | 6J9E (109) | RNAP ω | *Xanthomonos oryzae* | 8.6 | 3.7 |

**Fig. 6.**
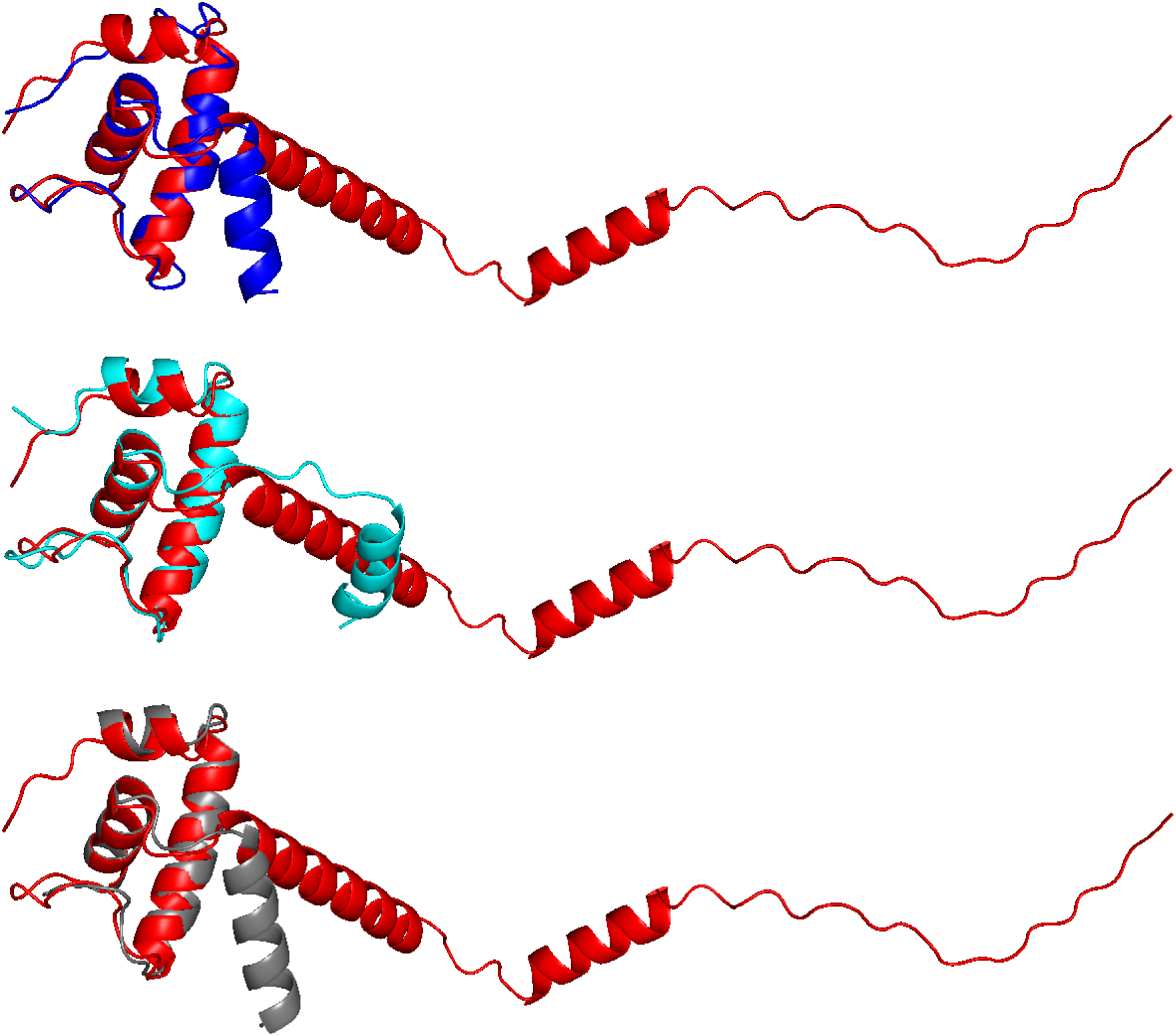
AlphaFold predictions for annotated ω of *Candidatus Midichloria* RNAP ω. Superimposition of AlphaFold prediction for Candidatus Midichloria RNAP ω (red) on experimental structures of *Pseudomonas aeruginosa, M. tuberculosis*, and *Xanthomonas oryzae* RNAP ω (blue, cyan, and gray, respectively).

### Absence of ω in *Mycoplasma* and *Ureaplasma*

We next extended our RNAP ω subunit search in the facultative intracellular bacterium *Mycoplasma genitalium*, whose 580-kb genome is the smallest known bacterial genome (110). NCBI search failed to identify an annotated *rpoZ* in *M. genitalium*. Interestingly, our search also failed to identify an annotated *rpoZ* in other *Mycoplasma* species, even though most have genome sizes comparable to that of *Chlamydia*. Our search also failed to identify an annotated *rpoZ* in *Ureaplasma* (111), which is phylogenetically closely related to *Mycoplasma*. To verify the absence of ω subunits in these organisms, we first checked the gene immediately downstream of the *gmk* gene in *Mycoplasma* for possible sequence similarity to *rpoZ*, and we found none (110, 111). We next performed AlphaFold modeling for all 68 hypothetical proteins of *Mycoplasma pneumoniae* having sizes comparable to bacterial ω subunits (i.e., sizes of 40-150 amino acids) (112). AlphaFold predicted multi-α-helix folds for 26 of the 68 proteins. Three-dimensional-structure similarity searches of these 26 AlphaFold predictions, performed on the DALI server (90, 91), failed to identify structures of experimentally determined bacterial ω subunits as possible matches. We infer that *Mycoplasma* and *Ureaplasma* are unlikely to have RNAP ω subunits.

## Discussion

In this report, we present multiple lines of evidence for the existence of an RNAP ω subunit in chlamydiae. Although a lack of strong, continuous sequence homology previously had precluded the identification of a chlamydial ω, a multi-step BlastP analysis led to the identification of CTL0286 as candidate (Fig. 1). Like *rpoZ* in the super majority of bacteria, *ctl0286* is located immediately downstream of *gmk* (Table 2 and data not shown). AlphaFold-predicted three-dimensional structures of CTL0286 exhibit strong similarities to experimental three-dimensional structures of ω subunits for a broad range of bacterial taxa (29, 101, 109, 113, 114). AlphaFold-Multimer predicted three-dimensional structures of complexes of CTL0286, with *C. trachomatis* RNAP β’ subunit, and of CTL0286 with *C. trachomatis* RNAP β’ and β subunits, exhibit strong similarity to experimental three-dimensional structures of ω-β’ and β’-β complexes [(Fig. 4, 5); (97, 98)]. The identification of CTL0286 as the *C. trachomatis* ω demonstrates the power of use of combinations of sequence-similarity analysis, synteny analysis, and AlphaFold and AlphaFold-Multimer analysis for identifying proteins “missing” from proteomes and for annotating functions of hypothetical proteins in proteomes.

Our extended analysis further showed that like *Chlamydia*, other obligate intracellular bacteria (i.e., *Rickettsia, Anaplasma, Ehrlichia, Orientia, Wolbachia* and *Candidatus Midichloria*) also encode ω orthologs (Fig. 6 and data not shown), but facultative intracellular bacteria *Mycoplasma* and *Ureaplasma* do not. Together with previous findings demonstrating the existence of ω orthologs in archaea and eukaryotes (27-29), these findings suggest that all living organisms from bacteria to humans have omega orthologs, likely with *Mycoplasma* and *Ureaplasma* as only exceptions.

ω plays roles in σ-RNAP core enzyme association (21-23) and thereby influences promoter-recognition selectivity (21-23). *Chlamydiae* possess a principal σ factor and two alternative σ factors (80, 81, 115). The principal σfactor, σ^66^, is involved in transcription of most chlamydial genes throughout the developmental cycle; the alternative σ factors, σ^28^ and σ^54^, are required for expression of certain late genes (116-118). The different chlamydial σ factors also differentially affect response to stress conditions (71, 77). It would be equally interesting to investigate if and how the chlamydial ω regulates σ-RNAP core enzyme association in chlamydial developmental stages and in response to various stress condition.

In summary, we have identified the long-missing ω subunit of the cRNAP. As with most scientific studies, this discovery raises more questions than it answers. There is a need to determine whether the chlamydial ω plays solely a structural role in cRNAP assembly and stability, or whether it also functions in regulation of chlamydial growth, development, and stress response.

## ACKNOWLEDGEMENTS

This work was supported by grants from the National Institutes of Health (AI071954 to HF and GM041376 to RHE). We thank Yu Zhang and Liqiang Shen for bringing *gmk-rpoZ* synteny to our attention and Jason Kaelber for helpful discussions.

